# Molecular basis of a dominant SARS-CoV-2 Spike-derived epitope presented by HLA-A*02:01 recognised by a public TCR

**DOI:** 10.1101/2021.08.15.456333

**Authors:** Christopher Szeto, Andrea T. Nguyen, Christian A. Lobos, Dimitra S.M. Chatzileontiadou, Dhilshan Jaya-singhe, Emma J. Grant, Alan Riboldi-Tunnicliffe, Corey Smith, Stephanie Gras

## Abstract

The data currently available on how the immune system recognizes the SARS-CoV-2 virus is growing rapidly. While there are structures of some SARS-CoV-2 proteins in complex with antibodies, which helps us understand how the immune system is able to recognise this new virus, we are lacking data on how T cells are able to recognize this virus. T cells, especially the cytotoxic CD8+ T cells, are critical for viral recognition and clearance. Here we report the X-ray crystallography structure of a T cell receptor, shared among unrelated individuals (public TCR) in complex with a dominant spike-derived CD8+ T cell epitope (YLQ peptide). We show that YLQ activates a polyfunctional CD8+ T cell response in COVID-19 recovered patients. We detail the molecular basis for the shared TCR gene usage observed in HLA-A*02:01+ individuals, providing an understanding of TCR recognition towards a SARS-CoV-2 epitope. Interestingly, the YLQ peptide conformation did not change upon TCR binding, facilitating the high-affinity interaction observed.

## 1. Introduction

Severe acute respiratory syndrome coronavirus 2 (SARS-CoV-2) is an emerging virus which has infected over 200 million people worldwide resulting in coronavirus disease 2019 (COVID-19), and over 4.3 million deaths (1). Despite the rapid development of effective and safe vaccinations against COVID-19, the global infection rate remains high, likely due to mutations within the SARS-CoV-2 virus, driven by the scale of global infections, and now vaccination, which pressures the virus to select for viral mutations that facilitate immune escape. Cytotoxic T cells are vital in the control and clearance of viral infections (2-5) and have been shown to be an important factor of the immune response to SARS-CoV-2, due their role in viral clearance and ability to recognize variants of SARS-CoV-2 (6). CD8+ T cells typically recognize peptides of 8-10 amino acid long presented by human leukocyte antigen (HLA) molecules (7).

To date, over 1000 distinct CD8+ T cell epitopes have been reported (www.iedb.org (8)), spanning multiple SARS-CoV-2 proteins. These epitopes are restricted by a large range of HLA class I (HLA-I) molecules, including HLA-A*02:01, one of the most prevalent HLAs in the global population (9). Several studies have shown that HLA-A*02:01+ individuals demonstrate a strong CD8+ T cell response to one such HLA-A*02:01 restricted CD8+ T cell epitope derived from the Spike (S) protein of SARS-CoV-2, namely, S_269–277_ (YLQPRTFLL, hereafter refer as YLQ) (10-18), which was characterised as an immunodominant epitope (18).

CD8+ T cells recognise the peptide-HLA complex (pHLA) though their T cell receptor (TCR). TCRs comprise an α- and β-chain, composed of variable (V), joining (J), constant (C) and diversity (D; β-chain only), genes generated by somatic recombination (19). Additional diversity is introduced by the inclusion of non-template encoded (N) regions at the junction of gene segments by the terminal deoxynucleotidyl transferase (tdt) enzyme, leading to incredible diversity (20). Indeed, it is estimated that there are 2×10^7^ TCR combinations available in humans (21). Within the TCR, three regions of variability, termed complementarity determining regions (CDRs) exist and are responsible for TCR specificity (7). Of these, the CDR3 region, which spans the V(D)J gene segments, is the most variable (7), and has been shown both functionally and structurally to make the predominant contacts within the pHLA complex (7).

The CDR3αβ loops are typically used to define peptide-specific CD8+ T cell clonotypes and the combination of these clonotypes is referred to as the TCR repertoire. TCR repertoires can exhibit biases, that is a preference for particular TCR α-chain Variable (TRAV) or TCR β-chain Variable (TRBV) usage (7, 22, 23). Additionally, despite the vast array of potential TCRs in any given individual, identical epitope-specific clonotypes have been described across donors. These “public” TCRs are thought to have a selective advantage, or comprise predominately germline encoded sequences that could be easily generated in different individuals (7, 23-25). However, TCR repertoires are more typically private, where each individual displays completely distinct TCR sequences specific for the same epitope (26-28). Understanding the TCR repertoire, and in the case of public TCRs, how they interact with their pHLA molecule, is critical for a thorough understanding of CD8+ T cell response towards specific epitopes.

Here, we wanted to validate and dissect the CD8+ T cell response to the YLQ peptide and determine the structural basis for the presentation of the YLQ peptide by HLA-A*02:01. Additionally, we aimed to provide the molecular basis of the biased TCR repertoire observed in response to the YLQ epitope in COVID-19 recovered individuals in different studies (16-18). Therefore, we have selected a representative public TCR, hereafter called YLQ-SG3 TCR. We determined the ternary structure of the HLA-A*02:01-YLQ peptide bound to the public YLQ-SG3 TCR and investigated the binding affinity of the public TCR.

## 2. Materials and Methods

### Sequence alignment

The full spike proteins from the five different coronaviruses were aligned using the online alignment software Rhône-Alpes Bioinformatics Center (PRABI http://www.prabi.fr/) multiple sequence alignment CLUSTALW (29). The accession number for the sequence used were for SARS-CoV-2: YP_009724390.1, OC43: YP_009555241.1, HKU-1: AZS52618.1, 229E: AAG48592.1, NL63: AAS58177.1. Then the sequence aligned with the SARS-CoV-2 YLQ peptide was selected and reported in **Table 1**.

**Table 1.** YLQ homologues peptides from seasonal coronaviruses

| Virus | YLQ homologue | Sequence identity (%) |
| --- | --- | --- |
| SARS-CoV-2 | YLQPRTFLL | - |
| OC43 | PLTSRQYLL | 44 |
| HKU-1 | PLSRRQYLL | 44 |
| 229E | ALPKTVREF | 11 |
| NL63 | FGPSSQPYY | 0 |

### SARS-CoV-2 YLQ conservation

The sequence conservation of the YLQ peptide was obtained using the NCBI web site tool “Mutations in SARS-CoV-2 SRA Data” (https://www.ncbi.nlm.nih.gov/labs/virus/vssi/#/scov2_snp) that uses the Wuhan-Hu-1 as reference sequence of SARS-CoV-2 with the accession number of NC_045512.2. The web site was accessed on the 30^th^ of July 2021, with 412,297 full sequences of the spike proteins were available.

### Generation of peptide-specific CD8+ T cell lines

CD8+ T cell lines were generated as previously described (26, 30). In summary, HLA typed HLA-A*02:01+ PBMCs from COVID-19 recovered individuals were incubated with SARS-CoV-2 peptide pools (2µM / peptide) and cultured for 10-14 days in RPMI-1640 supplemented with 1x Non-essential amino acids (NEAA; Sigma), 5 mM HEPES (Sigma), 2 mM L-glutamine (Sigma), 1x penicillin/streptomycin/Glutamine (Life Technologies), 50 µM 2-ME (Sigma) and 10% heat-inactivated (FCS; Thermofisher, Scientifix). Cultures were supplemented with 10IU IL-2 (BD Biosciences) 2-3 times weekly. CD8+ T cell lines were freshly harvested and used for subsequent assays.

### Intracellular cytokine assay

The intracellular cytokine assay was performed as previously described (26, 30). Briefly, CD8+ T cell lines were stimulated with cognate peptide pools or 10µM individual peptides (Genscript) and incubated for 5 hours in the presence of GolgiPlug (BD Biosciences), GolgiStop (BD Biosciences) and anti-CD107a-AF488 (BD Biosciences/eBioscience). Following incubation, cells were surface stained for 30 minutes with anti-CD8-PerCP-Cy5.5 (eBioscience/BD Biosciences), anti-CD4-BUV395 (BD Biosciences) anti-CD14-APCH7, CD19-APCH7 and Live/Dead Fixable Near-IR Dead Cell Stain (Life Technologies). Cells were then fixed and permeabilized for 20 minutes using BD Cytofix/Cytoperm solution (BD Biosciences) and intra-cellularly stained with anti-IFN-γ-BV421, and anti-TNF-PE-Cy7(all BD Biosciences) for a further 30 minutes. Cells were acquired on a BD LSRFortessa with FACSDiva software. Analysis was performed using FlowJo software where cytokine levels identified in the R0 control condition were subtracted from corresponding test conditions.

### Protein refold, purification, crystallisation

The HLA-A*02:01 heavy chain and β2-microglobulin as well as both chains of the YLQ-SG3 TCR were produced using bacterial expression of inclusion bodies and refolded into soluble protein (for detailed protocol see (31)). In brief, DNA plasmids encoding each recombinant protein subunit (HLA-A*02:01 α-chain, β2-microglobulin, TCR α-chain, and TCR β-chain) were individually transformed into competent BL21 *E. coli* cells. All cells were grown separately and their inclusion bodies were extracted. Soluble HLA-A*02:01-YLQ complex was produced by refolding inclusion bodies in the following amounts: 30 mg of α-chain, 10 mg of β2-microglobulin and 4 mg of YLQ peptide (Genscript). Soluble YLQ-SG3 TCR was produced by refolding 50 mg of TCRα chain with 50 mg of TCRβ chain. The refold buffer used was 3 M Urea, 0.5 M L-Arginine, 0.1 M Tris-HCl pH 8.0, 2.5 mM EDTA pH 8.0, 5 mM glutathione (reduced), 1.25 mM glutathione (oxidised). The refold mixtures were separately dialysed into 10 mM Tris-HCl pH 8.0. HLA-A*02:01-YLQ was purified using anion exchange chromatography (HiTrap Q, GE), whilst the YLQ-SG3 TCR was purified using anion exchange followed by size exclusion chromatography (Superdex 200 16/60, GE).

Crystals of HLA-A*02:01-YLQ complex were obtained using the sitting-drop, vapour-diffusion method at 20 °C with a protein/mother liquor drop ratio of 1:1 at 6 mg/mL in 10 mM Tris-HCl pH 8.0, 150 mM NaCl using 20% PEG3350 and 0.2 M NaF. YLQ-SG3 TCR was co-complexed with HLA-A*02:01-YLQ by combining both proteins at a 1:1 molar ratio before purification using size exclusion chromatography (Superdex 200 10/30, GE). Crystals of YLQ-SG3 TCR-HLA-A*02:01-YLQ complex were obtained using the sitting-drop, vapour-diffusion method at 20°C with a protein/mother liquor drop ratio of 1:1 at 3 mg/mL in 10 mM Tris-HCl pH 8.0, 150 mM NaCl using 20% PEG3350 and 0.05 M Zn-Acetate. Crystals were soaked in a cryosolution of 30% (w/v) PEG3350 diluted using mother liquor and then flash frozen in liquid nitrogen. The data were collected on the MX2 beamline at the Australian Synchrotron, part of ANSTO, Australia (32).

### Structure determination

The data were processed using XDS (33) and the structures were determined by molecular replacement using the PHASER program (34) from the CCP4 suite (35) using a model of HLA-A*02:01 without peptide (derived from PDB ID: 3GSO (36)). Manual model building was conducted using COOT (37) followed by refinement with BUSTER (38). The final models have been validated and deposited using the wwPDB OneDep System and the final refinement statistics, PDB codes are summarized in **Table 3**. All molecular graphics representations were created using PyMOL (Schrodinger, LLC, v1.7.6.3).

### Stability assay

Thermal stability was measured using differential scanning fluorimetry, performed in a Qiagen RG6 rtPCR. HLA-A*02:01-YLQ was heated from 30 to 95°C at a rate of 0.5°C/min with excitation and emission channels set at yellow (excitation of ∼530 nm and detection at ∼557 nm). The experiment was performed at two concentrations (5 µM and 10 µM) in duplicate. Each sample was dialysed in 10 mM Tris-HCl pH 8.0, 150 mM NaCl and contained a final concentration of 10X SYPRO Orange Dye. Fluorescence intensity data was normalised and plotted using GraphPad Prism 9 (version 9.0.0).

### Surface Plasmon resonance (SPR)

SPR was performed using a Biacore T200 biosensor at 25°C. YLQ-SG3 TCR was immobilized onto a CM5 chip using amine coupling, with the reference flow cell containing a negative control (M1_58-66_ TCR (23)). The immobilization steps were carried out at a flow rate of 5 µl/min in immobilization buffer 10 mM HEPES (pH 7.0), 150 mM NaCl and finally blocked with Ethanolamine at 5 µl/min for 7 min. HLA-A*02:01-YLQ was injected over the chip at a range of concentrations from 0.2 to 50 µM using a 1 in 2 dilution at a flow rate of 30 µl/min and in a running buffer of 10 mM Tris–HCl pH 8.0, 150 mM NaCl, 1 mg/ml bovine serum albumin and 0.005% P20. All injections were run in duplicate and SPR was performed twice to determine the dissociation constant between YLQ-SG3 TCR and HLA-A*02:01-YLQ (n=2) using both steady state affinity measurements and kinetics data. Kinetics data was analysed using the T200 BiaE-valuation software, whilst steady state values were extracted using T200 BiaEvaluation software, plotted and fitted into a one-site specific binding non-linear regression using Graphpad Prism (version 9.0).

## 3. Results

### 3.1. The YLQ epitope induced a polyfunctional CD8^+^ T cell response in COVID-19 recovered donors

The CD8+ T cell response towards the HLA-A*02:01 restricted YLQ peptide has previously been reported (10, 11, 16-18), however data regarding the level of polyfunctionality associated with the CD8+ T cell response has been limited in COVID-19 recovered donors. Therefore, we first tested the immunogenicity of the YLQ peptide in three COVID-19 recovered individuals by expanding CD8+ T cells against peptide pools including the YLQ peptide and performed an intracellular cytokine staining assay to determine the immunogenicity. The CD8+ T cell response and cytokine production towards the YLQ peptide was variable between the COVID-19 recovered donors. CD8+ T cells from two out of three donors, namely Q036 and Q042, were able to produce all four cytokines, while only double cytokine producing CD8+ T cells were observed in the Q062 donor (**Figure 1**). Even though the level of polyfunctionality was different between the three donors, they were all able to generate a polyfunctional CD8+ T cell response specific to the YLQ peptide after recovery from COVID-19.

**Figure 1.**
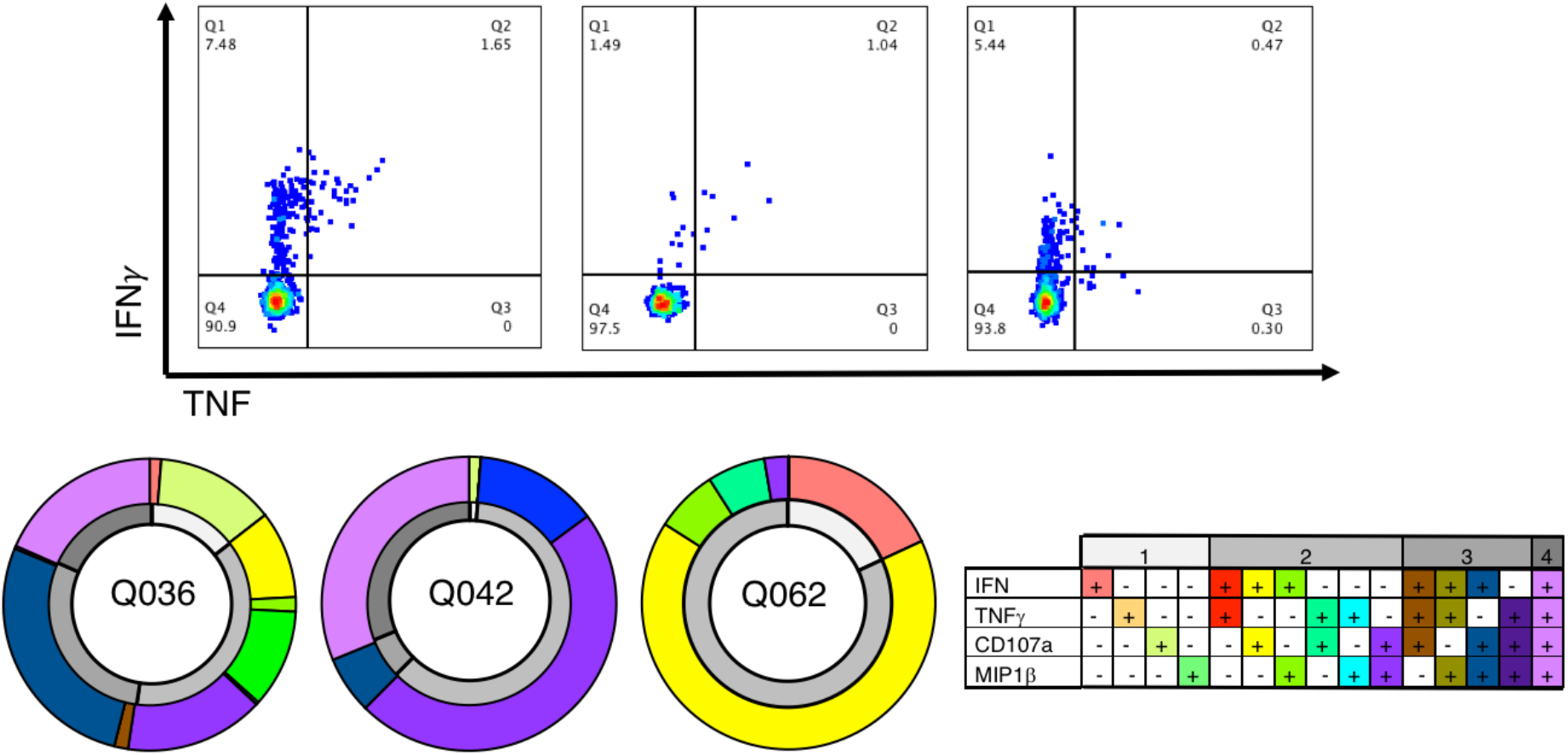
Polyfunctional response from YLQ-specific CD8+ T cells. The top panels are representative FACs plot for each donor (Q036, Q042, and Q062) showing IFNγ and TNF production from CD8+ T cells against the YLQ peptide. The bottom panels represent the polyfunctionality level of CD8+ T cells for the same COVID-19 recovered donors as the top panel. The outside ring, in colour, shows the % of cytokine combinations produced by CD8+ T cells, while the inner ring (in grey) represent the number of functions produced by CD8+ T cells. The key for each colours are in the bottom right table.

### 3.2. The conserved YLQ peptide is stably presented by the HLA-A*02:01 molecule

We previously determined that a high level of CD8+ T cell activation towards a SARS-CoV-2 epitope derived from the nucleocapsid (N_105-113_ or SPR peptide) was underpinned by a pre-existing and cross-reactive response (26). This preexisting immunity was due to a high level of sequence identity (55-89%) between the SPR peptide from SARS-CoV-2 and its homologues from seasonal coronaviruses. Therefore, we questioned if the YLQ peptide was also conserved in seasonal coronaviruses by aligning the spike protein sequences of SARS-CoV-2 and seasonal coronaviruses (**Table 1**).

While the SPR peptide had up to 89% sequence identity with its seasonal coronaviruses derived homologues, the level of conservation of the YLQ peptide was lower. The YLQ peptide shared only four residues with its homologues from OC43 and HKU-1 β-coronaviruses, with two of those residues being primary anchor residues that will be buried in the HLA cleft. The low sequence identity is in line with the lack or weak T cell activation observed in healthy individuals (18).

As YLQ peptide is derived from spike protein, which is relatively less conserved and more prone to mutation than other SARS-CoV-2 viral proteins, we also wanted to assess the level of mutations found in the different SARS-CoV-2 isolates (**Table 2**). Interestingly, this dominant T cell epitope was conserved with less than 0.5% of mutations for any of its residues.

**Table 2.** YLQ conservation in SARS-CoV-2 isolates (non-synonymous mutation)

| Residue | Y | L | Q | P | R | T | F | L | L |
| --- | --- | --- | --- | --- | --- | --- | --- | --- | --- |
| mutation | H<br>(0.006%) | V<br>(0.005%) | K<br>(0.005%) | S<br>(0.012%) | K<br>(0.014%) | S<br>(0.004%) | L<br>(0.002%) |  |  |
|  | D<br>(0.005%) |  | E<br>(0.003%) | L<br>(0.431%) | M<br>(0.009%) | I<br>(0.014%) |  |  |  |
|  | C<br>(0.005%) |  | R<br>(0.009%) | H<br>(0.012%) | S<br>(0.021%) |  |  |  |  |
|  |  |  | L<br>(0.002%) |  |  |  |  |  |  |
| % variant | 0.016 | 0.005 | 0.021 | 0.456 | 0.045 | 0.018 | 0.002 | 0 | 0 |

In order to gain a deeper understanding of the YLQ peptide recognition by CD8+ T cells, we first refolded and crystallised the HLA-A*02:01-YLQ complex and solved its structure at a high resolution (**Table 3**). The electron density map was clear for the peptide, indicating a stable and rigid conformation of the YLQ peptide in the HLA-A*02:01 cleft (**Figure 2A-B**).

**Table 3.**
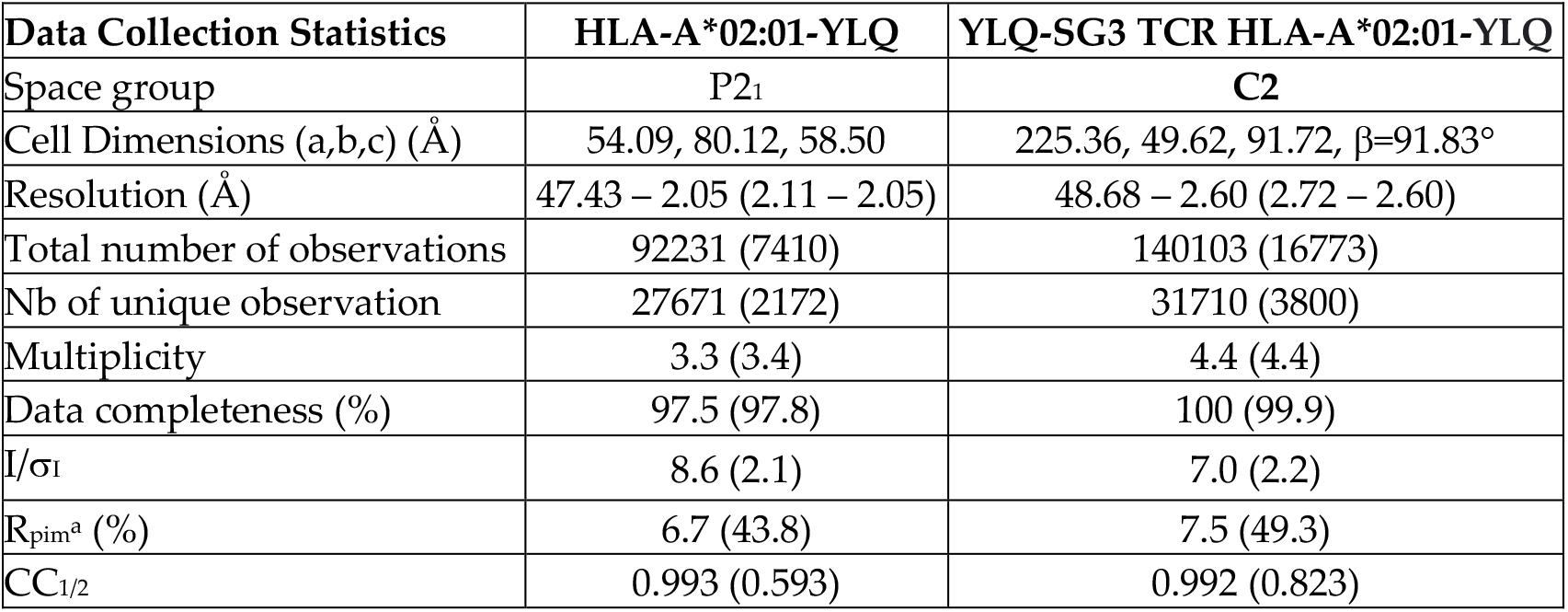
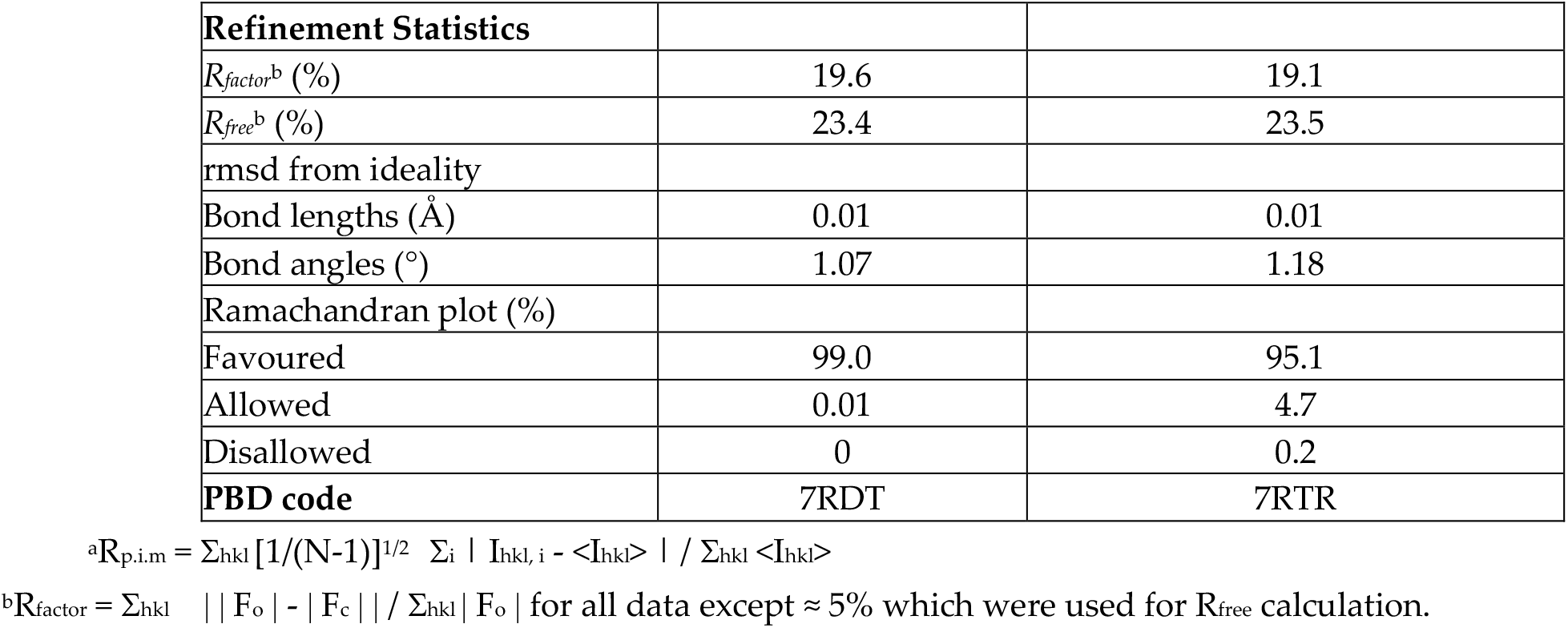
Data Collection and Refinement Statistics

**Figure 2.**
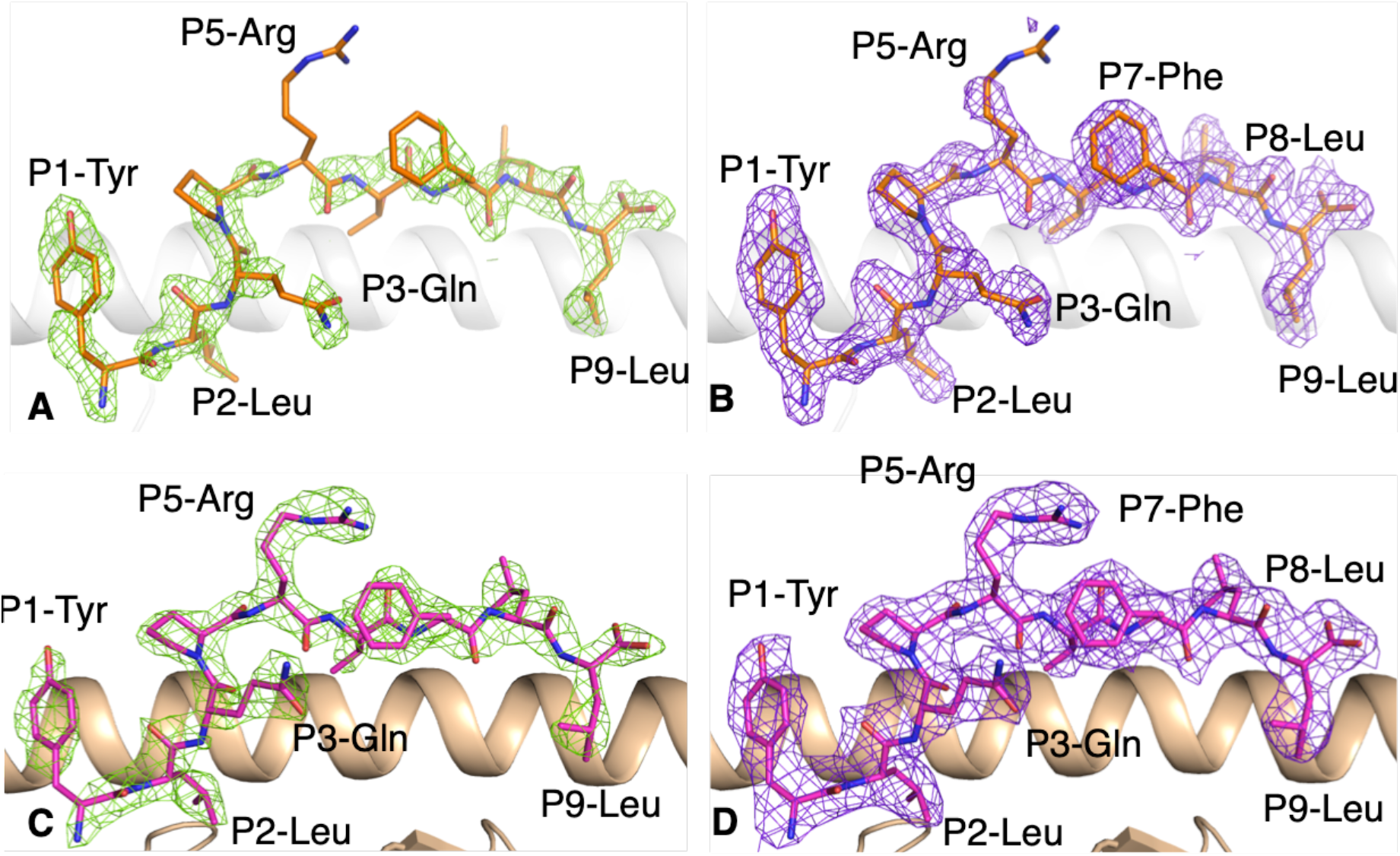
Electron density map for the YLQ peptide bound to HLA-A*02:01 without and with the YLQ-SG3 TCR. (**A, B**) Electron density map of (**A**) Fo-Fc map at 3σ (green) and (**B**) 2Fo-Fc at 1 σ (purple) around the YLQ peptide (orange stick) in complex with HLA-A*02 :01 (white cartoon). (**C, D**) Electron density maps of (**C**) Fo-Fc map at 3σ (green) and (**D**) 2Fo-Fc at 1 σ (purple) around the YLQ peptide (pink stick) presented by the HLA-A*02:01 (beige cartoon) bound to the YLQ-SG3 TCR.

The YLQ peptide bound to the HLA-A*02 :01 cleft via the canonical primary anchor of small hydrophobic residues, P2-Leu and P9-Leu, characteristic of HLA-A*02 :01, and an additional secondary anchor with P3-Gln (**Figure 2A-B**). The YLQ peptide has a series of residues with long, solvent exposed side-chains, P1-Tyr, P5-Arg, P7-Phe and P8-Leu, that could potenially interact with TCRs. The side-chains were well defined in the electron density map, this is possibly due to the numerous intra-peptide contacts. The only exception was the P5-Arg for which the density was partly missing for the side-chain, showing high mobility (**Figure 2B**). This rigidity of the pHLA was apparent when we undertook a thermal stability assay to determine the stability of the overall peptide-HLA (pHLA) complex, as this is important for immunogenicity (30). Indeed, the thermal stability of the HLA-A*02:01-YLQ complex was about 60°C, which is similar to that observed for the dominant influenza derived M1_58-66_ peptide bound to HLA-A*02:01 (23, 30).

### 3.3. The dominant YLQ peptide is recognised by public TCRs

The YLQ peptide was reported to be immunogenic in ∼90% of COVID-19 recovered individuals, while only 5% of healthy donors exhibited T cells specific for the peptide (18). This shows that in the absence of an antibody response, this epitope can be used as a marker of infection in HLA-A*02:01+ patients, and also that a T cell driven immune response would be activated. Interestingly, three studies have reported the TCR sequences of YLQ-specific clonotypes from COVID-19 recovered individuals and show an highly biased repertoire among unrelated donors (**Table 4**).

**Table 4.**
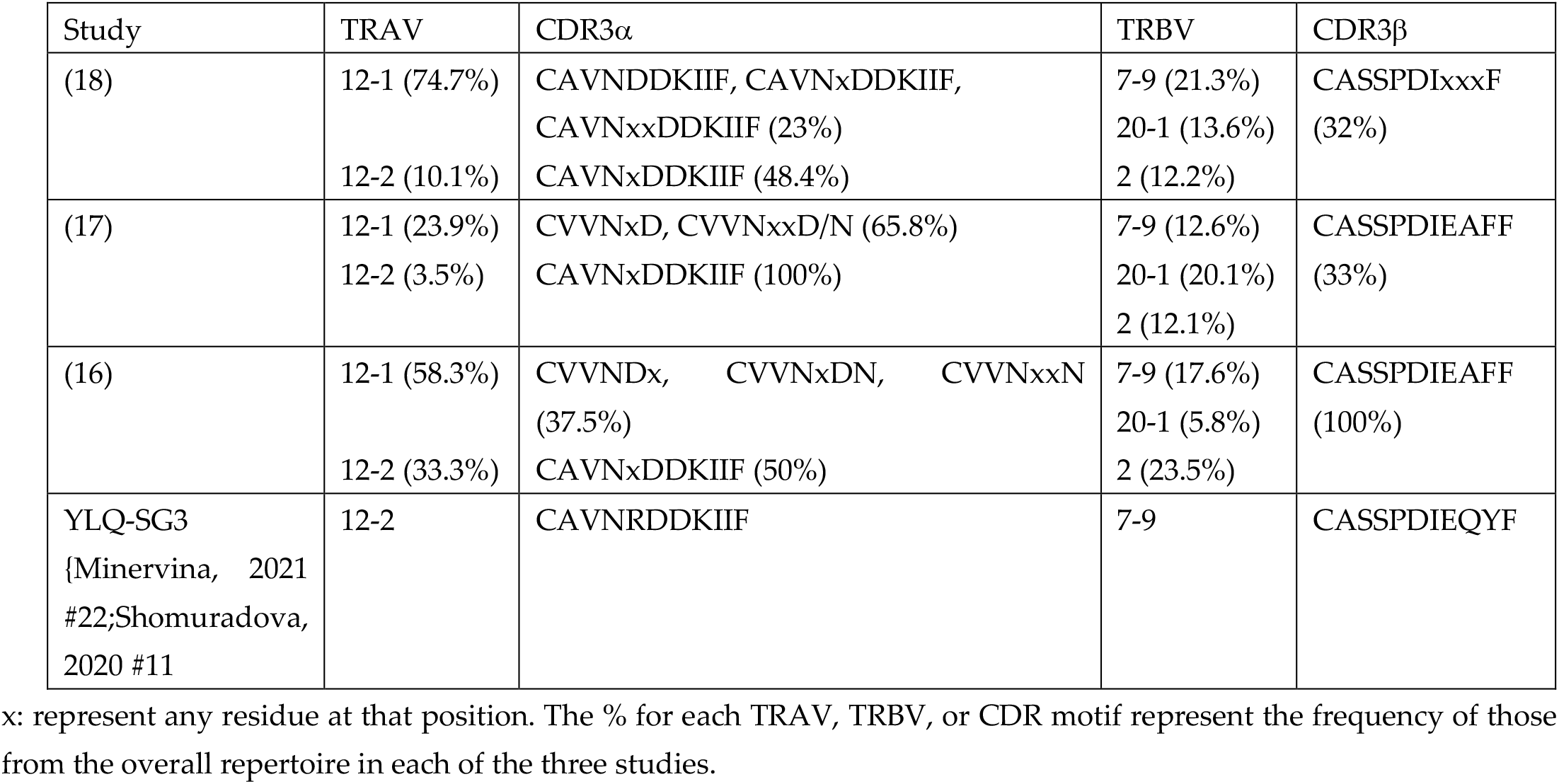
YLQ-specific biased TCR repertoire

We analysed the TCR sequences from those studies, and observed the same TCR gene usage bias, especially for the TCR α-chain. The HLA-A*02:01-YLQ-specific T cells were mostly expressing a TRAV12-1 or TRAV12-2 allele for their α-chain, both sharing 50% sequence identity for their CDR1α and CDR2α loops. The most frequent TRBV gene expressed by YLQ-specific CD8+ T cells were 2, 7-9, and 20-1 with different frequencies depending on the study (**Table 4**). Interestingly, there were conserved motifs present in both α and β CDR3 loops, with a public TCR observed among donors and across studies, here called the YLQ-SG3 TCR (**Table 4**). The YLQ-SG3 TCR was composed of the TRAV12-2 and TRBV7-9 bias chain and contains the conserved motif within both its CDR3 loops. We therefore chose the YLQ-SG3 TCR to understand how T cells can engage with YLQ epitope, a SARS-CoV-2 spike-derived peptide presented by the HLA-A*02:01 molecule.

### 3.4. Structure of the public YLQ-SG3 TCR recognising the dominant YLQ epitope presented by HLA-A*02:01

We refolded and purified the YLQ-SG3 TCR and undertook affinity measurements by surface plasmon resonance (SPR), as well as solved the structure of the YLQ-SG3 TCR in complex with the HLA-A*02:01-YLQ.

The SPR data shows that the YLQ-SG3 TCR binds with the HLA-A*02:01-YLQ complex with high affinity and a Kd of 2.09 ± 0.16 µM (**Figure 3A-B**), at the high end of the affinity range observed for CD8+ TCR (7). In addition the kinetics of the interaction show a fast association (k_on_ = 386,800 ± 25,000 M^-1^s^-1^) and a fast dissociation (koff = 0.679 ± 0.001 s^-1^) compare to other TCRs (27) (**Figure 3A**).

**Figure 3.**
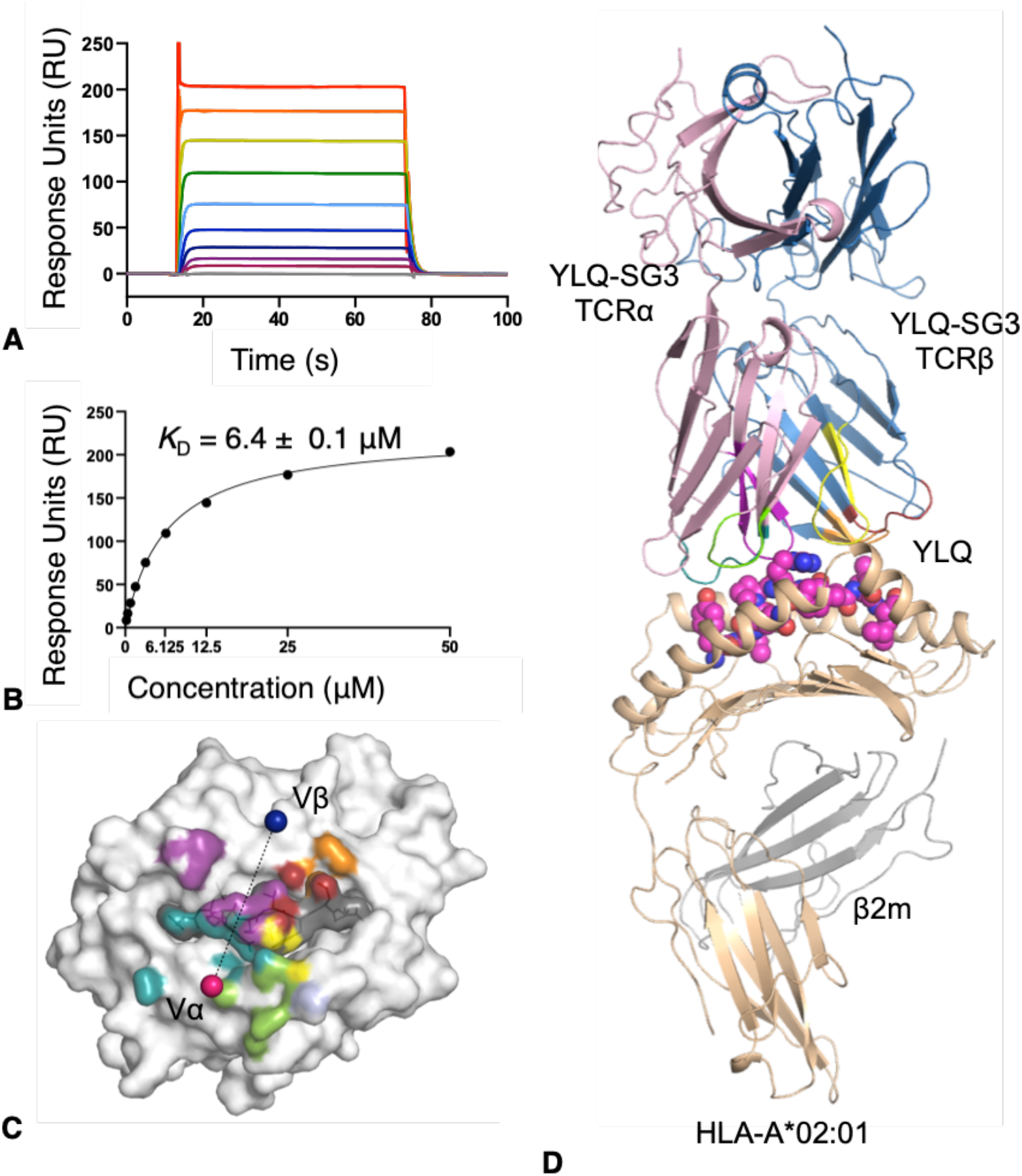
YLQ-SG3 TCR affinity and structure with HLA-A*02:01-YLQ. (**A**) SPR sensorgram and (**B**) steady state binding curve for YLQ-SG3 TCR towards HLA-A*02:01-YLQ. Analyte HLA-A*02:01-YLQ was flowed over immobilized YLQ-SG3 TCR with a concentration range of 0.19 to 50 μM. (**C**) Footprint of the YLQ-SG3 TCR shows atomic contact with the HLA-A*02:01-YLQ complex. The surface of HLA-A*02:01 is in white, the surface of the YLQ peptide is in grey. Each atoms are colored by the TCR segment they are contacted by, with for the CDR1/2/3α in deepteal, chartreuse and purple, for the CDR1/2/3β in red, orange and yellow, and β-chain framework in light blue. The pink and blue spheres represent the mass center of the α- and β-chain, respectively. (**D**) overview of the YLQ-SG3 TCR (α-chain in pink, β-chain in blue) represented as cartoon on the top of the YLQ peptide (pink spheres) presented by the HLA-A*02:01 (beige cartoon, with the β2m in grey cartoon). The TCR CDR loops are colored as per panel C.

We solved the structure of the YLQ-SG3 TCR in complex with the HLA-A*02:01-YLQ to better undertsand how TCRs recognise SARS-CoV-2 epitope. We solved the structure at a resolution of 2.6 Å (**Table 3**) with unambigous density for the peptide (**Figure 2C-D**).

The YLQ-SG3 TCR docks diagonally above the center of the YLQ peptide with a docking angle of 73° (**Figure 3B-C**), within the range of other TCR-pHLA complexes (7). The buried surface area at the interface of the TCR and HLA-A*02:01-YLQ was 1,809 Å, also within the range (average of 1,885 Å) (7). Interestingly, and consistent with the strong TCR bias observed for the YLQ-specific T cells, the TCR α-chain is contributing to 67% of the interaction (**Figure 3B**), with the CDR1/2α loops contributing to 40% of the total interactions and giving the molecular basis for the TRAV12 bias observed (**Table 4**). All CDRα loops contacted the HLA-A*02:01 molecule, however, from these, only CDR1α and CDR3α contacted the YLQ peptide (**Table 5**).

**Table 5.** Contacts between the YLQ-SG3 TCR and HLA-A*02:01-YLQ complex.

| TCR segment | TCR residues | HLA-A*02:01 residues | Type bond |
| --- | --- | --- | --- |
| CDR1α | Arg28<sup>Nη1-Nε</sup> | Glu166<sup>Oε2-Oε1</sup> | VDW, SB |
| CDR1α | Gln37<sup>Nε2</sup> | Gln155<sup>O</sup>, Tyr159 | VDW, HB |
| CDR1α | Ser38<sup>Oγ</sup> | Gln155<sup>Oε1</sup> | VDW, HB |
| FWα | Phe55 | His151 | VDW |
| CDR2α | Tyr57 | Glu154, Gln155, Ala158 | VDW |
| CDR2α | Ser58<sup>Oγ</sup> | Glu154<sup>Oε2</sup>, Arg157<sup>Nη2</sup> | VDW, HB |
| CDR3α | Asp109<sup>Oδ1-Oδ2</sup> | Arg65<sup>Nε-Nη2</sup>, Lys66 | VDW, SB |
| CDR1β | Arg38 | Thr73 | VDW, HB |
| CDR2β | Gln57<sup>Nε2</sup> | Thr73<sup>Oγ1</sup>, Val76 | VDW |
| CDR2β | Asn58 | Val76 | VDW |
| CDR3β | Asp109<sup>Oδ2</sup> | Ala150, Gln155<sup>Nε2</sup> | VDW, HB |
| CDR3β | Ile110 | Gln115 | VDW |
| TCR segment | TCR residues | YLQ peptide residues | Type bond |
| CDR1α | Asp27<sup>Oδ1</sup> | Tyr1<sup>O$H$</sup> | HB |
| CDR1α | Gly29 | Tyr1, Pro4 | VDW |
| CDR1α | Gln37<sup>Oδ1</sup> | Gln3<sup>Nε2</sup>, Pro4, Arg5 | VDW, HB |
| CDR1α | Ser38 | Arg5 | VDW |
| CDR3α | Asn107 | Arg5 | VDW |
| CDR3α | Asp109<sup>O-Oδ1</sup> | Pro4, Arg5<sup>Nε-Nη1</sup> | VDW, HB, SB |
| CDR3α | Asp110<sup>Oδ1-Oδ2</sup> | Arg5, Thr6<sup>O-N-Oγ</sup> | VDW, HB |
| CDR1β | Asn37 | Leu8 | VDW |
| CDR1β | Arg38<sup>Nη2</sup> | Arg5, Thr6<sup>O</sup>, Leu8 | VDW, HB |
| CDR2β | Gln57 | Leu8 | VDW |
| CDR3β | Asp109<sup>O-Oδ2-Oδ1</sup> | Arg5<sup>Nη1-Nη2</sup>, Leu8 | VDW, HB, SB |
| CDR3β | Ile110 | Arg5 | VDW |

Abbreviations are as follows: FW, framework residue; HB, hydrogen bond (cut-off distance 3.5Å); SB, salt bridge (cut-off distance 5Å); VDW, van der Waals (cut-off distance 4 Å).

The CDR1α loop streched itself above the N-terminal region of the α2-helix and forms a salt bridge with Glu166 via Arg28α, as well as hydrogen bonds with the Gln155 via the Gln37α and Ser38α. In addition, the side-chain of the Gln37α dips in between the HLA α2-helix and the peptide backbone to form an extensive hydrogen bond network (**Figure 4A**). The CDR2α sits above the α2-helix of the HLA just before the hinge region of the α2-helix, with the Ser58α forming H-bond with Arg157 outside the cleft, and the Tyr57 forming Van der Waals bonds with the Gln155 inside the cleft (**Figure 4B**). The CDR3α makes limited contributions in forming contacts with the HLA molecule, with the conserved Asp109α (**Table 5**) forming a salt bridge with the Arg65 (**Figure 4C**).

**Figure 4.**
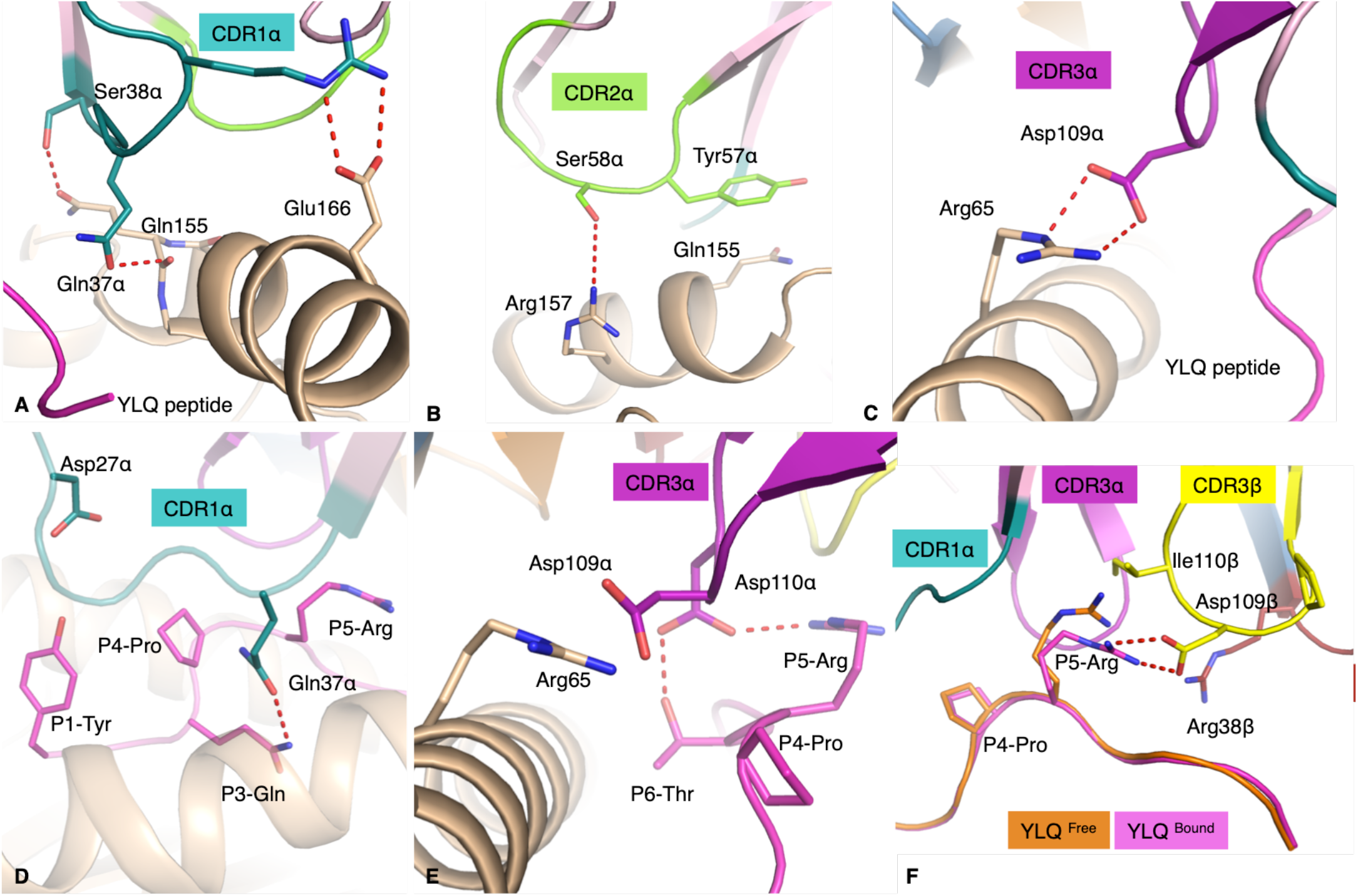
YLQ-SG3 TCR interaction with the HLA-A*02:01 molecule and the YLQ peptide. All the panels show the interaction between the YLQ-SG3 TCR with the α-cahin in pale pink; the β-chain in pale blue; the CDR1/2/3α loops in deepteal, chartreuse and purple, respectively; and the CDR1/2/3β coloure in red, organe and yellow, respectively. The HLA-A*02:01 is coloured in beige, and the YLQ peptide either in pink when bound to the YLQ-SG3 TCR or orange if free. The red dashed lines represent hydrogen bonds. (**A**) CDR1α (deepteal) interacting with the α2-helix of the HLA-A*002:01, with the YLQ peptide in pink cartoon. (**B**) CDR2α (chartreuse) inteacting with the α2-helix of the HLA before the hinge region of the molecule. (**C**) CDR3α loop forming a salt bridge with the HLA Arg65 residue, the YLQ peptide is in pink cartoon. (**D**) CDR1α Gln37α inserting its side-chain in between the HLA (beige cartoon) and the peptide backbone (pink) maximising the surface interaction with the peptide. (**E**) CDR3α forming a network of hydrogen bonds with the YLQ peptide. (**F**) Superposition of the YLQ peptide structures with (pink) and without (orange) the YLQ-SG3 TCR, also showing the interaction of the CDR3β (yellow) with the peptide (pink).

The YLQ-SG3 TCR β-chain has limited contact with the HLA. Both CDR1β and CDR2β loops made contacts with two residues of the α1-helix and the CDR3β loop with two residues of the α2-helix (**Table 5**).

The YLQ peptide made a significant contribution to the pHLA buried surface area at 38% and is contacted by five of the CDR loops (**Figure 3C** and **Table5**), whilst the average buried surface area is only 29% for other solved TCRpMHC complexes (7). The CDR1α loop runs over half of the peptide making contacts from P1-Tyr to P5-Arg, with the side-chain of the Gln37α inserting itself between the peptide and α2-helix and interacting with P3-Gln, P4-Pro and P5-Arg (**Figure 4D**). In the same fashion, the CDR3α loop contacts a large strecth of the YLQ peptide including P4-Pro, P5-Arg and P6-Thr, and inserts a conserved CDR3α^109^DD^110^ motif in between the peptide and the α1-helix of the HLA-A*02:01 (**Figure 4E**). The CDR1β and CDR2β loops each projected long side-chains towards the C-terminal parts of the peptide surface. As as result the exposed P8-Leu is surrounded by Asn37β/Arg38β on one side and by Gln57β/Asn58β on the other side. The CDR3β pushes the P5-Arg down with the Ile110β and forms a salt bridge with the Asp109β. This conformation is helped by the short length of the CDR3β loop that only forms a short rigid loop due to the Pro108β. The P5-Arg is surrounded by the CDR1/3α and CDR1/3β loops (**Figure 4F**), and instead of wrapping the side-chain of the P5-Arg with CDR loops that has been previously observed (39), the YLQ-SG3 TCR pushes down on the P5-Arg and P6-Phe side-chains. This increases the contact surface between the peptide and these loops and stabilised the P5-Arg side-chain, yet, do not disturb the HLA-A*02:01 cleft structure (root mean square deviation of 0.22 Å). Overall the YLQ-SG3 TCR docks onto HLA-A*02:01-YLQ with minimal structural rearrangements, with the exception of a few residue side-chains. As the kinetics data from SPR shows a fast association rate (**Figure 3A**), high binding affinity and moderate dissociation rate. This is consistent with the larger binding interface, but minimal structural rearrangements during binding.

## 4. Discussion

We have described here the molecular basis of a public TCR recognizing a dominant spike-derived SARS-CoV-2 epitope. The structure of the YLQ peptide in the cleft of the HLA-A*02:01 molecule is a constrained and rigid peptide that forms numerous intra-peptide interactions favoured by large side-chain residues. This rigidity was consistent with the high thermal stability observed for the HLA-A*02:01-YLQ complex. This rigid conformation of the YLQ peptide did not undergo large structural changes, beside the stabilisation of the P5-Arg, upon the YLQ-SG3 TCR docking. Despite a solvent exposed P5-Arg, the side-chain of this residue was pushed down by the CDR3β loop to maximize the contact between the TCR and the YLQ peptide. This resulted in a large contribution of the peptide to the pHLA surface buried area of 38%, well above the average of 29% (7), highlighting the importance of the peptide in driving the interaction with the YLQ-SG3 TCR.

The YLQ-specific T cells exhibited a bias in their TCR repertoire with frequent usage of the TRAV12 gene for the α-chain. Here, we show the molecular basis behind this bias, as the TCRpHLA complex structure shows that the α-chain dominates this interaction, contributing to 67% of the TCR contact surface. This was mainly due to a large footprint of the CDR1α (26%) on the peptide, CDR2α (14%) on the HLA, and CDR3α (19%) that binds both the peptide and the HLA.

Interestingly, TRAV12 usage in TCRs that recognise HLA-A*02:01 has been observed in 45% of the TCRpHLA-A*02:01 structures solved (17/38), with 18% (7/38) using the TRAV12-2*01 gene (7). While the TRAV12+ TCRs all used their α-chains to contact the N-terminal parts of the peptide and the HLA-A*02:01 cleft, their CDR1α loops don’t necessarily share the same interactions. For example, although the CDR1α loop of the CD8 (40) and 868 TCRs (41) interacts similarly to the CDR1α of YLQ-SG3 TCR, the DMF5 TCR uses its CDR1α loop mainly to contact the peptide N-terminal part (42), and not the HLA. Another example is the NYE_S1 TCR, which docks with a tilt that pushes the CDR1α loop away from the pHLA and does not make contact (43). This shows, that while some common interactions between the TRAV12+ TCRs and the HLA-A*02:01 are consistent between some TCRs, there is also a large variability of docking modes for the same αTCR segment.

The malleability of docking while using the same or similar sequence between different TCR chains, show that the conserved motif observed in the YLQ-specific TCRs, while not identical, could lead to the same mode of recognition of the YLQ epitope. The sequence differences between these TCRs could be important to give the TCR repertoire enough breadth to recognize variants of the YLQ peptide, which could be critical in recognizing emerging mutations located in this region of spike.

The YLQ epitope has been identified as one of the dominant CD8+ T cell epitopes in individuals expressing the HLA-A*02:01 allele. Information about the immune response to the YLQ epitope will be critical in understanding the potential of the YLQ peptide as a targetable epitope for T cell-based therapeutics, biomarkers or vaccines against COVID-19. Firstly, the YLQ peptide is highly immunogenic in most COVID-19 recovered HLA-A*02:01+ individuals, and weakly or not recognized in healthy individuals (18), so in absence of antibodies this could be used as a marker of infection. Secondly, none of the current mutations reported for that region of the spike are within the Variant of Concern (VOC) or of Interest (VOI). The YLQ epitope selects for biased and public TCRs (22) that could give a selective advantage to HLA-A*02:01+ individuals. The public TCR exhibits a high affinity within the range of other potent anti-viral CD8+ T cells (7). And finally, we report here that in COVID-19 recovered individuals there is a polyfunctional response from CD8+ T cell stimulated with the YLQ peptide. The ability of the YLQ epitope to strongly stimulate CD8+ T cells has also been observed in vaccinated individuals (44). Altogether this makes the YLQ peptide a promising target to prime and boost CD8+ T cells against SARS-CoV-2 infection.

## Supplementary Materials

**Figure S1.**
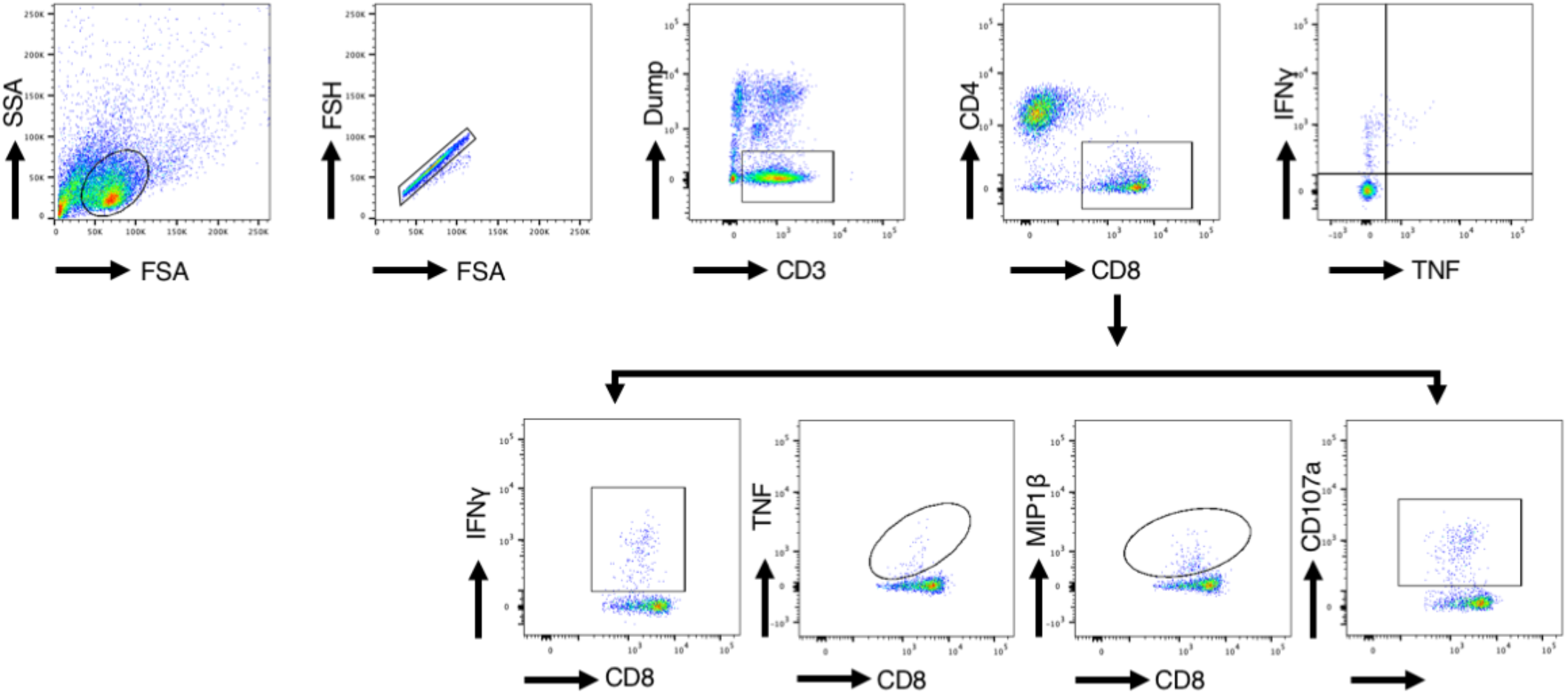
Gating strategy used in this study. (**A**) Gating strategy used to assess the functional responses of CD8+ T cell lines. Cells were gated on lymphocytes, singlets, CD3+Live cells, CD8+CD4-T cells to observe IFNγ and TNF production. (**B**) Gating strategy used to assess the polyfunctional responses of CD8+ T cell lines. Cells were gated on lymphocytes, singlets, CD3+Live cells, CD8+CD4-T cells and each of CD8+IFNγ+, CD8+TNF+, CD8+MIP1β and CD8+CD107a.

## Author Contributions

Conceptualization, S.G.; methodology, C.S. A.T.N., D.J. and S.G.; software, A.R.T. and C.S Szeto.; validation, C.S Szeto., A.T.N., and S.G.; formal analysis, C.S., A.T.N., and S.G.; investigation, C.S. A.T.N. and S.G.; resources, C.Szeto, C.Smith, and S.G.; data curation, C.L. and D.S.M.C..; writing—original draft preparation, S.G.; writing—review and editing, C.S Szeto., A.T.N., E.J.G. and S.G.; visualization, C.S Szeto., A.T.N., and S.G.; supervision, S.G.; project administration, S.G.; funding acquisition, C.Szeto, C.Smith, and S.G. All authors have read and agreed to the published version of the manuscript.

## Institutional Review Board Statement

This study was performed according to the principles of the Declaration of Helsinki. Ethics approval to undertake the research was obtained from the QIMR Berghofer Medical Research Institute Human Research Ethics Committee and Monash University Human Research Ethics Committee. COVID-19-recovered donors were over the age of 18, had been clinically diagnosed by PCR with SARS-CoV-2 infection, and had subsequently been released from isolation following resolution of symptomatic infection, and recruited in May and June 2020 from the south-east region of Queensland, Australia. The majority of participants were returned overseas travellers. Blood samples were collected from all participants to isolate peripheral blood mononuclear cells (PBMCs).

## Informed Consent Statement

Informed consent was obtained from all subjects involved in the study.

## Data Availability Statement

The final crystal structure models for the YLQ-HLA-A*02:01 and YLQ-SG3 TCR-HLA-A*02:01-YLQ complexes have been deposited to the Protein Data Bank (PDB) under the following accession codes: 7RDT and 7RTR, respectively.

## Acknowledgments

The authors would like to thank Queensland Health Forensic & Scientific Services, Queensland Department of Health who provided the SARS-CoV-2 isolate QLD02; Monash Macromolecular Crystallisation Facility; MX team for assistance at the Australian Synchrotron and the Australian Cancer Research Foundation (ACRF) Eiger detector. The authors would also like to thank all the participants who took part in the study.

